# Altered Capicua expression drives regional Purkinje neuron vulnerability through ion channel gene dysregulation in Spinocerebellar ataxia type 1

**DOI:** 10.1101/2020.05.21.104976

**Authors:** Ravi Chopra, David D Bushart, John P Cooper, Dhananjay Yellajoshyula, Logan M Morrison, Haoran Huang, Daniel R Scoles, Stefan M Pulst, Harry T Orr, Vikram G Shakkottai

**Author notes:** Contributed equally (Ravi Chopra, David D Bushart). Correspondence to: Vikram G. Shakkottai, 4009 BSRB, 109 Zina Pitcher Place, Ann Arbor, MI 48109.

## Abstract

Selective neuronal vulnerability in neurodegenerative disease is poorly understood. Using the ATXN1[82Q] model of spinocerebellar ataxia type 1 (SCA1), we explored the hypothesis that regional differences in Purkinje neuron degeneration could provide novel insights into selective vulnerability. ATXN1[82Q] Purkinje neurons from the anterior cerebellum were found to degenerate earlier than those from the nodular zone, and this early degeneration was associated with selective dysregulation of ion channel transcripts and altered Purkinje neuron spiking. Efforts to understand the basis for selective dysregulation of channel transcripts revealed modestly increased expression of the ATXN1 corepressor Capicua (Cic) in anterior cerebellar Purkinje neurons. Importantly, lentiviral overexpression of Cic in the nodular zone accelerated both aberrant Purkinje neuron spiking and neurodegeneration. These findings reinforce the central role for Cic in SCA1 cerebellar pathophysiology and suggest that only modest reductions in Cic are needed to have profound therapeutic impact in SCA1.

## Introduction

The polyglutamine disorders are a class of autosomal dominant neurodegenerative diseases caused by a glutamine-encoding CAG triplet repeat (polyglutamine) expansion within the protein coding sequence for their respective disease gene. These disorders are characterized by neurodegeneration in a restricted subset of neuron types despite widespread pathogenic protein expression, a phenomenon often referred to as “selective vulnerability.” The importance of understanding selective vulnerability as a strategy for uncovering fundamental neurodegenerative disease pathways is widely acknowledged (Fu, Hardy, & Duff, 2018; Saxena & Caroni, 2011), but mechanisms of selective vulnerability remains poorly understood.

Spinocerebellar ataxia type 1 (SCA1) is a polyglutamine disorder caused by CAG repeat expansion within the Ataxin-1 (*ATXN1*) gene (Orr et al., 1993). Like other polyglutamine ataxias, SCA1 is characterized by predominant involvement of cerebellar and brainstem neurons (Rüb et al., 2012) despite widespread CNS expression of ATXN1 protein (Servadio et al., 1995). Cerebellar Purkinje neuron degeneration is prominent in SCA1 (Koeppen, 2005), suggesting that Purkinje neurons are particularly vulnerable to expression of polyglutamine-expanded ATXN1. This conclusion is supported by a transgenic model of SCA1 with expression of ATXN1 with 82 CAG repeats (ATXN1[82Q]) restricted to Purkinje neurons, resulting in prominent Purkinje neuron degeneration and motor impairment (Burright et al., 1995). It is well-understood that dysregulation of gene expression is central to SCA1 pathogenesis (Klement et al., 1998; Rousseaux et al., 2018), but the key disease-associated genes underlying Purkinje neuron degeneration are not well understood.

Relative sparing of Purkinje neurons in lobule X of the ATXN1[82Q] model has been noted (Clark et al., 1997), despite the fact that ATXN1[82Q] expression is driven by a promoter expected to produce widespread Purkinje neuron expression (Vandaele et al., 1991). This finding suggests that Purkinje neurons in the anterior cerebellum are particularly vulnerable to ATXN1[82Q] expression. Recognizing that efforts to understand selective vulnerability are confounded by the numerous differences between affected and unaffected cell types, we propose that regional differences in Purkinje neuron degeneration could reveal novel pathways central to disease.

In the current study, we explored the hypothesis that there are specific disease-associated genes whose regional pattern of dysregulation matches the regional pattern of Purkinje neuron degeneration. We demonstrated robust confirmation of prior qualitative observations that Purkinje neuron degeneration is accelerated in the anterior cerebellum (lobules II-V) relative to the posterior cerebellum (lobules IX-X). Consistent with our hypothesis, we found that not all disease-associated genes are dysregulated in a manner that correlates with this degeneration pattern. Instead, only a subset of disease-associated ion channel genes is dysregulated exclusively in anterior Purkinje neurons, and selective anterior Purkinje neuron firing abnormalities are observed that can be well-explained by changes in the affected channels. We identified modestly higher expression of the ATXN1 co-repressor, Capicua (Cic), in the anterior cerebellum in association with this regional ion channel gene suppression. Finally, we found that viral overexpression of Cic in the nodular zone recapitulated the firing abnormalities seen in the anterior cerebellum and also accelerated nodular zone neurodegeneration. Our findings underscore the central role for Cic in SCA1 cerebellar pathophysiology (Rousseaux et al., 2018). These findings provide important insights regarding Purkinje neuron vulnerability in SCA1 and elevate these channels as important targets for the development of disease-modifying therapy.

## Materials and Methods

### Mice

All animal procedures were approved by the University of Michigan Committee on the Use and Care of Animals. ATXN1[82Q] transgenic mice (Burright et al., 1995) were maintained homozygous for the transgene on an FVB/NJ background (Jackson Labs). In all experiments involving ATXN1[82Q] transgenic mice, age- and sex-matched wild-type FVB/NJ mice were used as experimental controls. *Atxn1*^*154Q*^ knock-in mice (Watase et al., 2002) were maintained on a C57Bl6J background (Jackson labs) by crossing *Atxn1*^*154Q/2Q*^ males with *Atxn1*^*2Q/2Q*^ wild-type females. In all experiments involving *Atxn1*^*154Q*^ knock-in mice, *Atxn1*^*2Q/2Q*^ littermates were used as wild-type experimental controls.

### Electrophysiology

#### Solutions

Artificial CSF (aCSF) contained the following (in mM): 125 NaCl, 3.5 KCl, 26 NaHCO3, 1.25 NaH2PO4, 2 CaCl2, 1 MgCl2, and 10 glucose. For all recordings except for measurements of capacitance, pipettes were filled with internal recording solution containing the following (in mm): 119 K Gluconate, 2 Na gluconate, 6 NaCl, 2 MgCl2, 0.9 EGTA, 10 HEPES, 14 Tris-phosphocreatine, 4 MgATP, 0.3 tris-GTP, pH 7.3, osmolarity 290. For measurements of capacitance, pipettes were filled with internal recording solution containing the following (in mM): 140 CsCl, 2 MgCl2, 1 CaCl2, 10 EGTA, 10 HEPES, 4 Na2ATP, pH 7.3, osmolarity 287 mOsm.

#### Preparation of brain slices for electrophysiological recordings

Mice were anesthetized by isoflurane inhalation, decapitated, and the brains were submerged in pre-warmed (36°C) aCSF. Slices were prepared in aCSF held at 36°C on a VT1200 vibratome (Leica) while being continuously bubbled with carbogen (95% 02/5% CO2). Slices were prepared to a thickness of 300 µm. Once slices were obtained, they were incubated in continuously carbogen-bubbled aCSF for 45 minutes at 34°C. Slices were subsequently stored in continuously carbon-bubbled aCSF at room temperature until use.

#### Patch clamp recordings

Only slices from the cerebellar vermis were utilized for recordings, and slices were selected based on the ability to clearly visualize lobules II-X. Purkinje neurons from cerebellar lobules II-V or lobules IX-X were identified for patch clamp recordings using a 40x water-immersion objective and infrared differential interference contrast (IR-DIC) optics that were visualized using NIS Elements image analysis software (Nikon). Recordings were made 1-5 hours after slice preparation in a recording chamber that was continuously perfused with carbogen-bubbled aCSF at 34°C at a flow rate of 2-3 mLs/min. Borosillicate glass patch pipettes were pulled with resistances of 3-5 MΩ. Recordings were performed on one of two patch clamp rigs, with equipment and technical specifications below:

- Rig 1: Data were acquired using an Axon CV-7B headstage amplifier, Axon Multiclamp 700B amplifier, Digidata 1440A interface, and pClamp-10 software (MDS Analytical Technologies). All voltage data were acquired in the current-clamp mode with bridge balance compensation and filtered at 2 kHz.
- Rig 2: Data were acquired using an Axon CV-7B headstage amplifier, Axopatch 200B amplifier, Digidata 1440A interface, and p-Clamp-10 software (MDS Analytical Technologies). All voltage data were acquired in the fast current-clamp mode (Magistretti, Mantegazza, Guatteo, & Wanke, 1996) and filtered at 2 kHz.

For a given experiment, all data were acquired on the same rig to avoid experimental confounds arising from data acquisition. In all cases, acquired data were digitized at 100 kHz. Cells were rejected if the series resistance rose above 15 MΩ, with the majority of recordings having a series resistance of <10 MΩ. Cells were also rejected if the series resistance changed by >20% during recording. Voltage in this study is corrected for a liquid junction potential of +10 mV (Telgkamp & Raman, 2002).

#### Analysis of cell capacitance

Determination of Purkinje neuron membrane capacitance was performed using a well-established method for analysis of a two-compartment equivalent circuit representing a Purkinje neuron (Llano, Marty, Armstrong, & Konnerth, 1991), and has been described previously (Chopra, Bushart, & Shakkottai, 2018; Stoyas et al., 2020). Briefly, recordings in the anterior cerebellum and nodular zone were performed in the presence of 50 µM picrotoxin (cat. no. P1675, Sigma-Aldrich), and capacitative transients were obtained in voltage clamp mode using 1 second steps to −90 mV from a holding potential of −80 mV. Recordings were discarded if the measured input resistance was <100 MΩ. Voltage steps were performed ten times and the recorded currents were averaged and low-pass filtered at 5 kHz. The decay of the capacitative transient was fit using a two-exponential decay function described below (Equation 1):

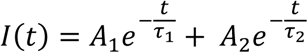

The constants from each cell’s decay function were then used to obtain four parameters: C_1_ (representing the capacitance of the soma and main proximal dendrites, Equation 2), C_2_ (representing the capacitance of the distal dendritic arbor, Equation 3), R_1_ (representing the pipette access resistance and internal resistance of the soma and proximal dendrite, Equation 4), and R_2_ (representing the composite internal resistance of dendritic segments separating the distal dendritic arbor from the main proximal dendritic segments, Equation 5). This was done as follows (equations described previously (Llano et al., 1991)):

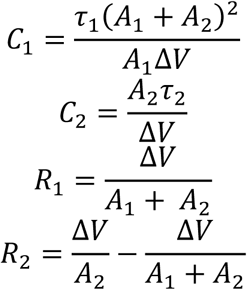

In our measurements of cell capacitance, |C_1_+C_2_| is reported in Figure 1B-1C.

**Figure 1.**
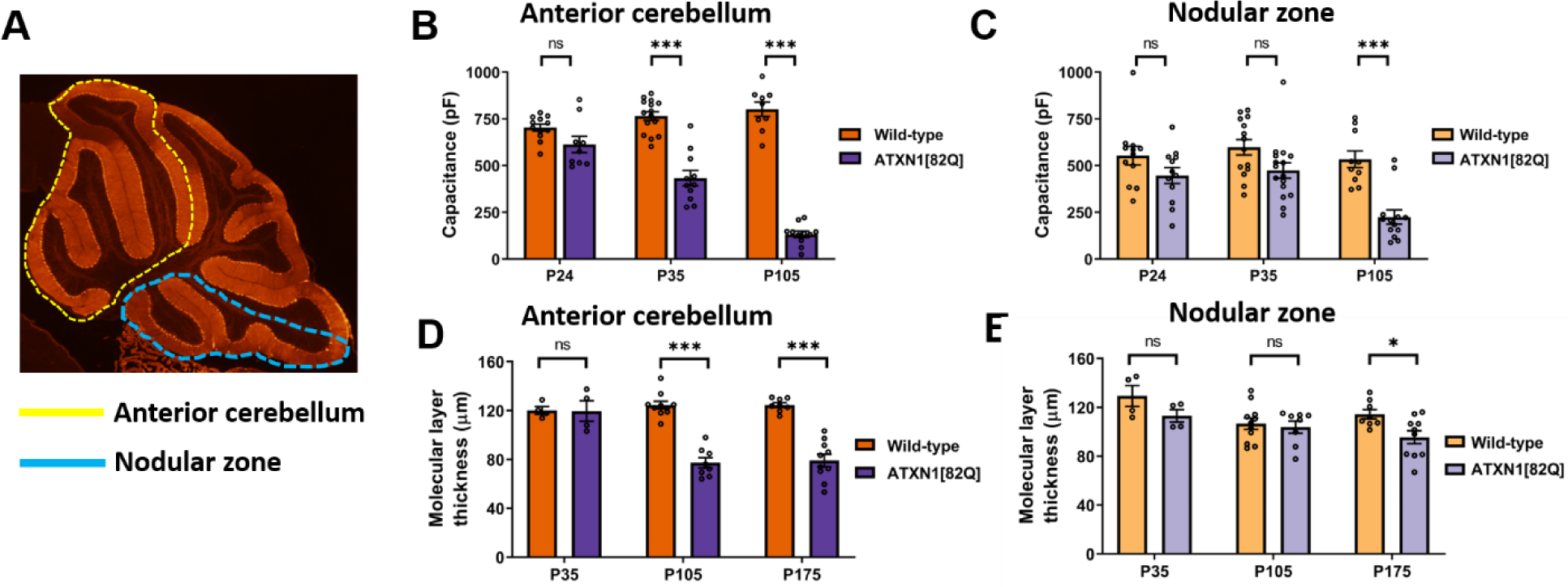
Purkinje neuron degeneration is delayed in the nodular zone of ATXN1[82Q] mice. (A) Diagram outlining the anterior cerebellar lobules (yellow dotted line) and nodular zone (blue dotted line). (B) Total Purkinje neuron capacitance was measured in the anterior cerebellar lobules of ATXN1[82Q] mice and wild-type controls at P24, P35, and P105. (C) Similar to panel (B), total Purkinje neuron capacitance was measured in the nodular zone at P24, P35, and P105. (D) Molecular layer thickness, a measurement which reflects the length of Purkinje neuron dendrites, was measured in the anterior cerebellar lobules of ATXN1[82Q] mice and wild-type controls at P35, P105, and P175. (E) Similar to panel (D), molecular layer thickness was measured in the nodular zone of ATXN1[82Q] mice and wild-type controls at P35, P105, and P175. * denotes p<0.05, *** denotes p<0.001, ns denotes p > 0.05; two way repeated-measures ANOVA with Holm-Sidak correction for multiple comparisons.

Of note, modeling studies have suggested that the reduced two-compartment Purkinje neuron model is statistically inadequate for estimating membrane capacitance in mature Purkinje neurons (Roth & Häusser, 2001). To ensure that our estimates based on the two-compartment model were internally consistent, these estimates were compared to capacitance estimates utilizing a different method wherein the capacitance is estimated from the area underneath the capacitative transient (Dell’Orco et al., 2015). Comparison of total cell capacitance by both methods revealed that there was no statistically-significant difference in estimated cell capacitance based on the method used, suggesting that the two compartment model could reliably estimate Purkinje neuron capacitance in our recordings (Supplementary Figure 1A-1D).

#### Analysis of Firing Properties

Electrophysiology data were analyzed offline using Clampfit 10.2 software (Molecular Devices). Firing frequency and coefficient of variation (CV) calculations were performed using a 10 s recording obtained in the cell-attached configuration ∼5 minutes after formation of a stable seal. The CV was calculated as follows:

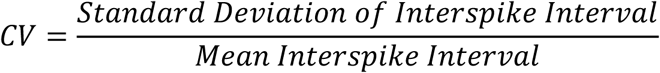

For Figures 3A, 3G, 3H, and 5B, cells were classified as either regular firing, irregular firing, or depolarization block/non-firing. For these Figures, irregular firing cells were defined as those cells whose CV was greater than the mean + 1 standard deviation CV of Purkinje neurons from the corresponding experimental control. Depolarization block/non-firing cells were defined as cells that were not firing in the cell-attached configuration but could fire a train of >5 action potentials in response to current injection in the whole-cell configuration. Cells which were not firing in the cell-attached configuration and were unable to fire a train of >5 action potentials in the whole-cell configuration were discarded.

**Figure 2.**
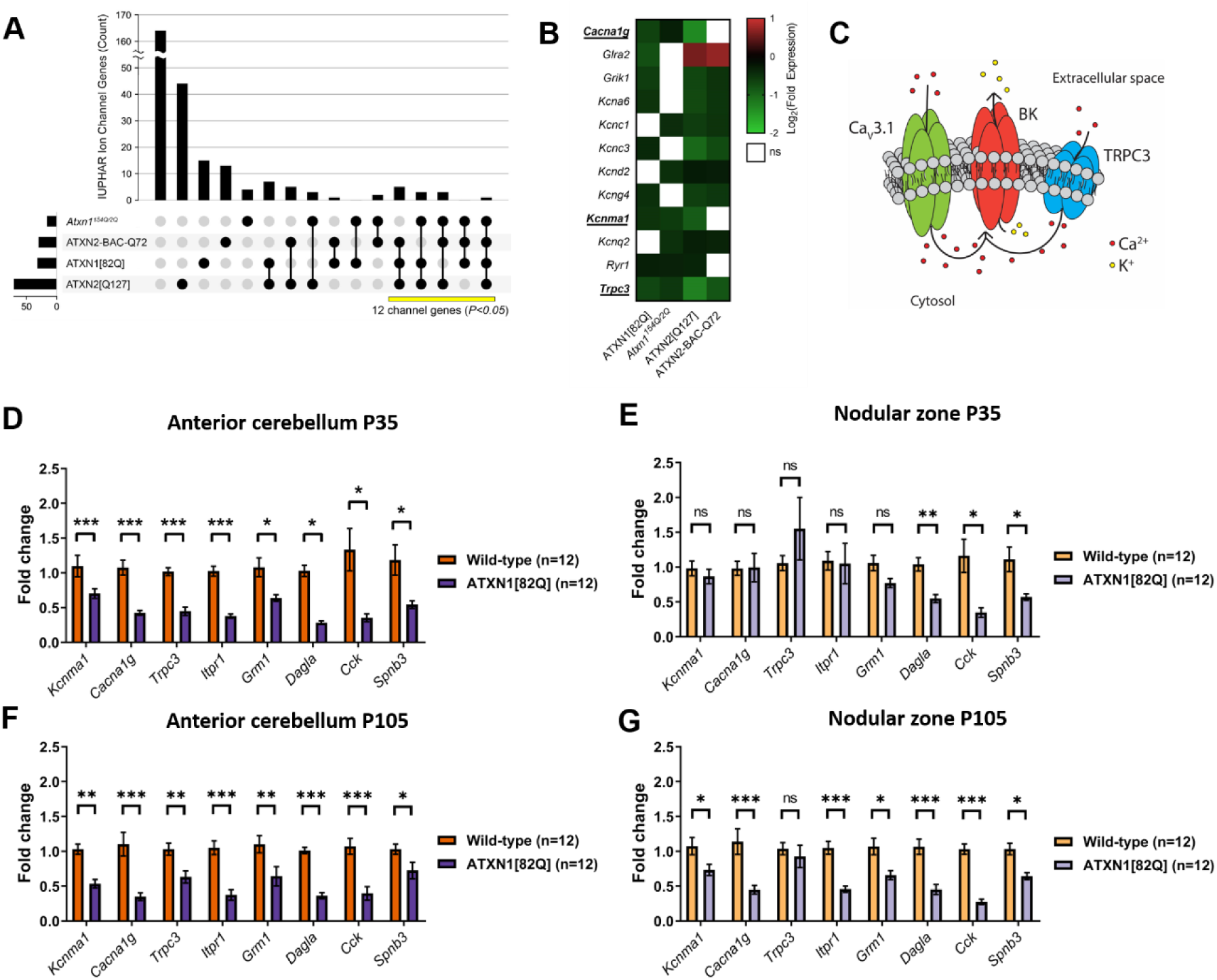
A functionally related module of channels is dysregulated uniquely in the anterior cerebellum of SCA1 mice. (A) IUPHAR-recognized ion channel genes that are dysregulated across models of SCA1 and SCA2. Vertical bars represent the number of ion channel genes unique to each overlap, and each overlap is delineated by black dots and linkages below the graph. The total number of ion channel genes differentially expressed in each model is shown as horizontal bars. The number of the channels that are found in at least three of four models (yellow bar) is statistically significant, with P-value reflecting the likelihood of an equivalent number of channels (or more) being dysregulated in these overlaps by chance (see methods section). (B) Twelve ion channel transcripts are among the genes that show downregulation across three or four models of SCA1 and SCA2. Three of these channel transcripts (*Cacna1g, Kcnma1*, and *Trpc3*) are known ataxia genes that form a putative functional ion channel module. (C) Proposed ion channel excitability module, highlighting a possible role for Ca_v_3.1 (encoded by *Cacna1g*) and TRPC3 (*Trpc3*) in providing Ca^2+^ for activation of BK channels (*Kcnma1*). (D-G) Quantitative real-time PCR (qRT-PCR) was performed in macrodissected cerebella from ATXN1[82Q] mice and wild-type controls at P35 from (D) anterior cerebellum or (E) nodular zone and at P105 from (F) anterior cerebellum or (G) nodular zone. The expression of all genes is represented as fold change (y-axis) with respect to wild-type. *denotes p<0.05; ** denotes p<0.01; *** denotes p<0.001; ns denotes p>0.05; two-tailed Student’s t-test with Holm-Sidak correction for multiple comparisons.

**Figure 3.**
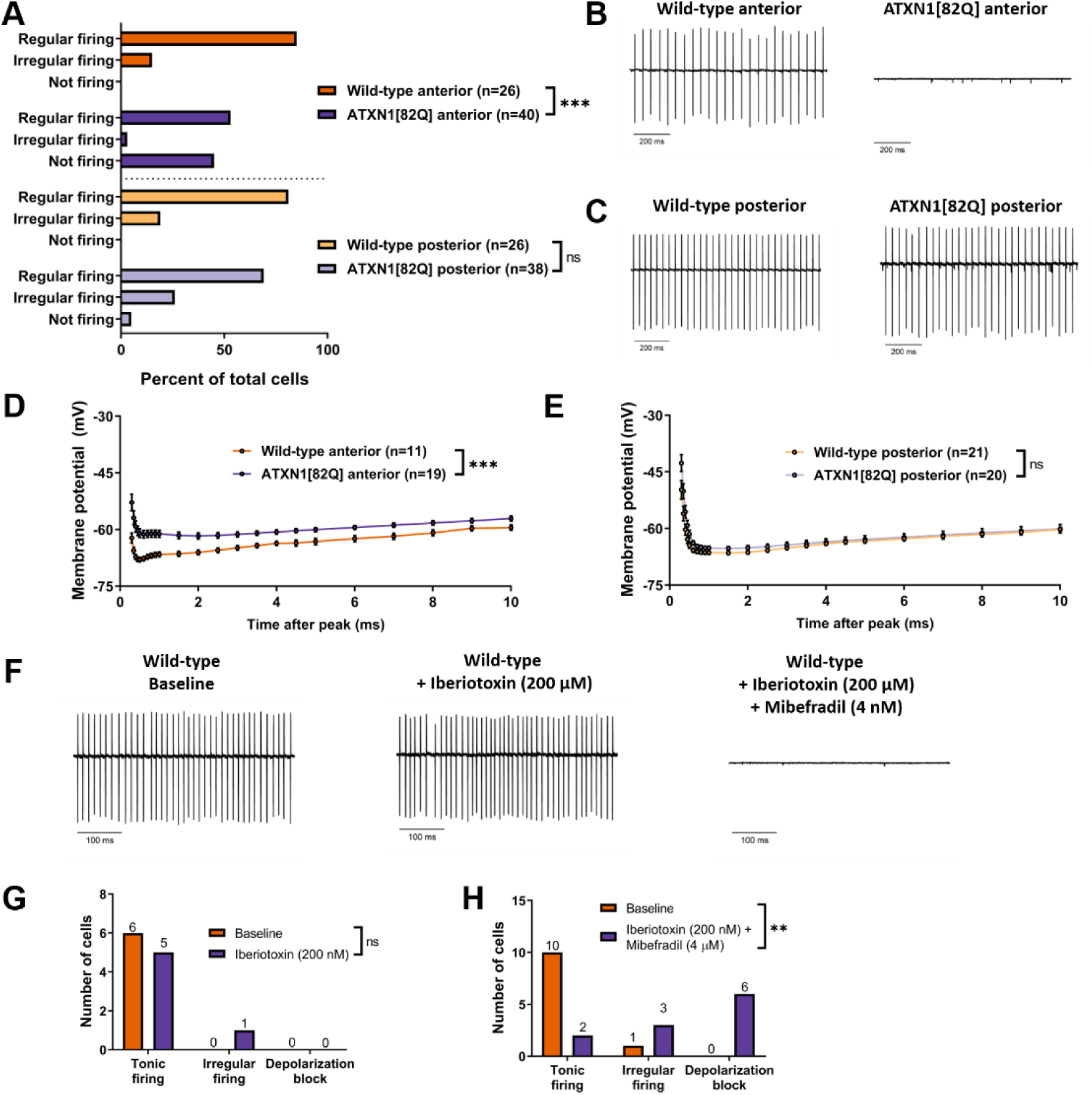
Regionally-dysregulated ion channel genes form a functional module critical for Purkinje neuron pacemaking. (A) The distribution of regularly firing, irregularly firing, and non-firing cells was recorded for Purkinje neurons in the anterior cerebellum and nodular zone. (B) Representative trace from a tonic firing wild-type and non-firing ATXN1[82Q] Purkinje neuron in the anterior cerebellum at P35. (C) Representative trace from wild-type and ATXN1[82Q] Purkinje neurons in the nodular zone at P35. (D) Quantification of the afterhyperpolarization (AHP) and ISI of wild-type and ATXN1[82Q] Purkinje neurons in the anterior cerebellum at P35. (E) Similar to (D), quantification of the AHP and ISI of wild-type and ATXN1[82Q] Purkinje neurons in the nodular zone at P35. (F) Representative traces of wild-type Purkinje neurons in the anterior cerebellum at P35. Traces are shown at baseline (left), after perfusion of 200 nM iberiotoxin (middle), and after perfusion of 200 nM iberiotoxin + 4 µM mibefradil. (G and H) Summary distribution of regularly firing, irregularly firing, and non-firing Purkinje neurons before and after perfusion of 200 nM iberiotoxin (G) and after 200 nM iberiotoxin + 4 µM mibefradil (H). ** denotes p<0.01; *** denotes p<0.001; ns denotes p>0.05; Chi-square test (A, G, H); two-way repeated measures ANOVA with Holm-Sidak correction for multiple comparisons (D, E).

**Figure 4.**
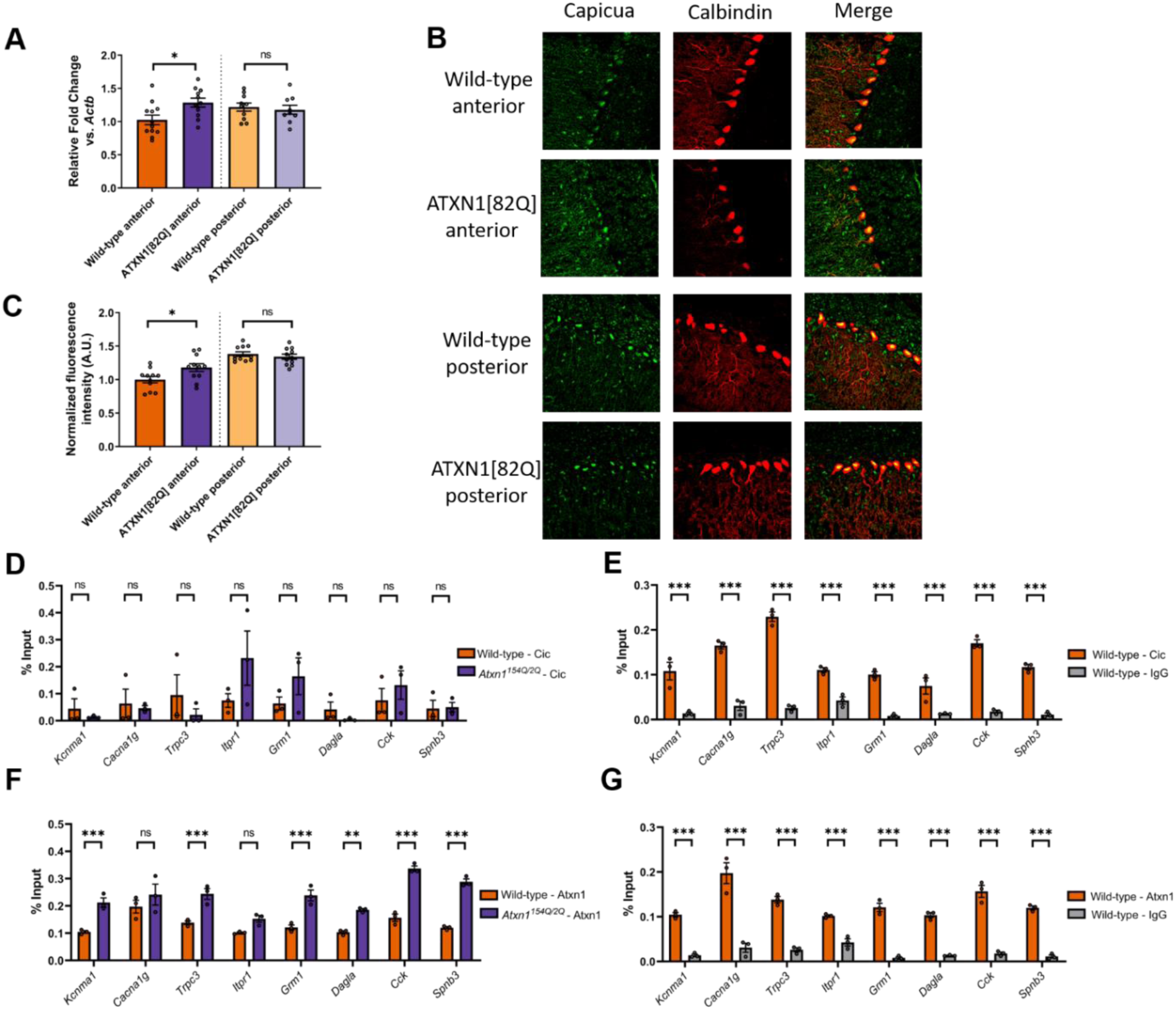
Higher Cic expression in Purkinje neurons from the anterior cerebellum of SCA1 mice is associated with repression of key ion channel genes. (A) Quantitative real-time PCR (qRT-PCR) was performed in the anterior cerebellum and nodular zone of ATXN1[82Q] mice and wild-type controls at P35. Relative fold change data compared to *Actb* are normalized to values from wild-type anterior cerebellum. (B) Representative confocal images taken from ATXN1[82Q] mice and wild-type controls at P35 after immunostaining for Capicua (green) and calbindin (red, to mark Purkinje neurons). (C) Quantification of relative expression of Capicua in the somata of ATXN1[82Q] and wild-type Purkinje neurons at P35, normalized to values from wild-type anterior cerebellum. (D-G) Quantitative Chromatin immunoprecipitation (qChIP) demonstrating the association of Capicua (Cic) and ATXN1 at the promoter of ion channel genes from sonicated chromatin derived from P14 whole cerebellar extracts. Binding, represented as % input (Y-axis) demonstrated for Cic (D-E) and ATXN1 (F-G) comparing their relative binding on ion channel genes in *Atxn1*^*154Q/2Q*^ mice and wild-type controls (D, F) and binding over background relative to their respective isotype control IgG (rabbit IgG) (E, G). * denotes p<0.05; ** denotes p<0.01; *** denotes p<0.001; ns denotes p>0.05; two-tailed Student’s t-test (A, C); two tailed-Student’s t-test with Holm-Sidak correction for multiple comparisons (D-I).

**Figure 5.**
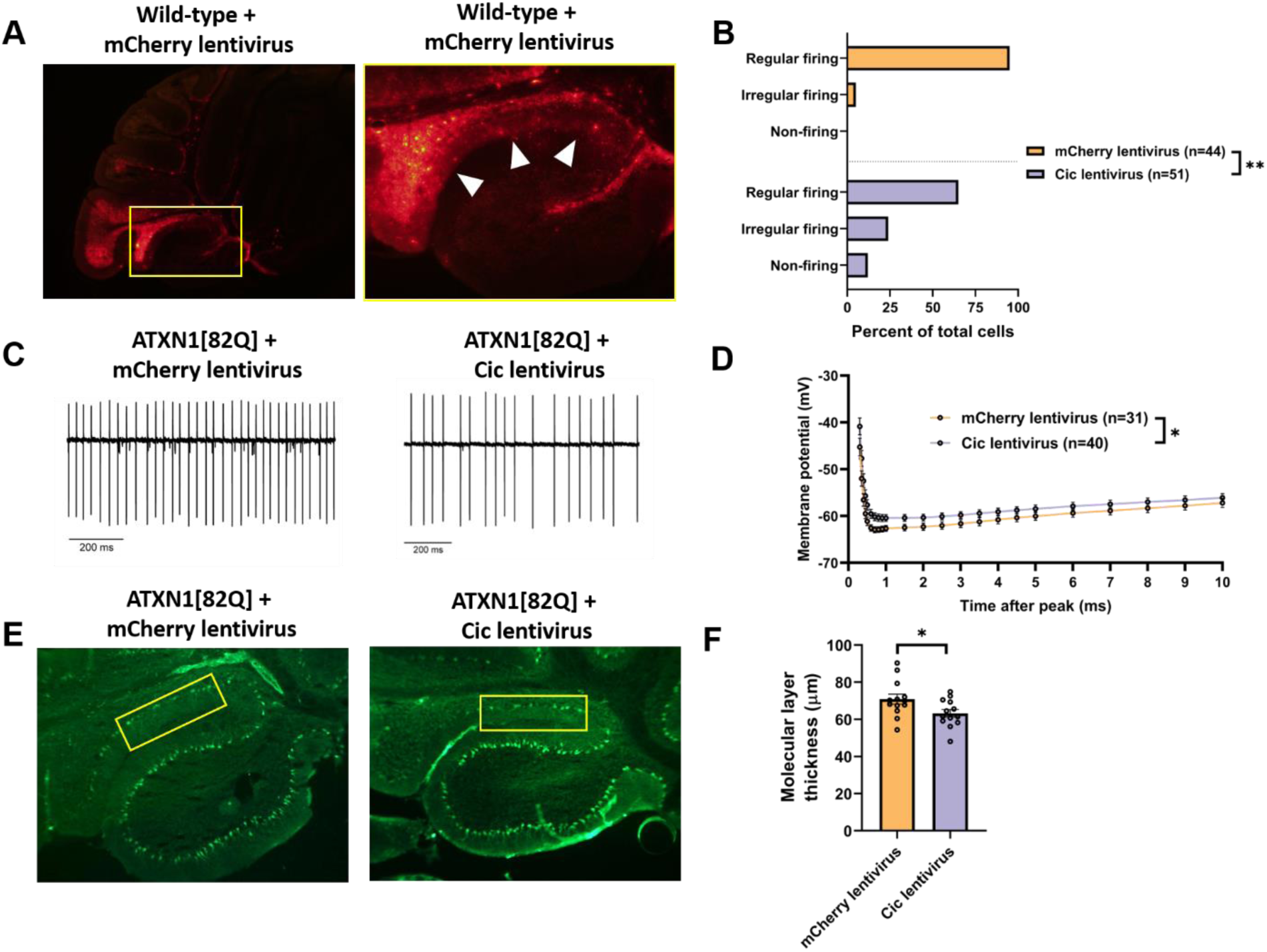
Increased Cic expression results in Purkinje neuron hyperexcitability and accelerates Purkinje neuron degeneration. (A) Stereotaxic injection of a lentiviral construct containing mCherry results in widespread expression in lobule IX, but not in lobule X, 10 days after injection. (B) The distribution of regularly firing, irregularly firing, and non-firing Purkinje neurons in the nodular zone of ATXN1[82Q] mice after 10 days of Cic lentivirus or control mCherry lentivirus expression. (C) Representative traces of ATXN1[82Q] Purkinje neuron spiking in the nodular zone 10 days after injection of Cic lentivirus or control mCherry lentivirus. (D) Quantification of the AHP from ATXN1[82Q] Purkinje neurons in the nodular zone, 10 days after injection of Cic lentivirus or control mCherry lentivirus. (E) Representative images of the nodular zone of ATXN1[82Q] mice at P105, 70 days post-injection of Cic lentivirus or mCherry control lentivirus. Yellow box indicates analyzed area of lobule IX. (F) Quantification of molecular layer thickness measurements as illustrated in panel (E). *denotes p<0.05; ** denotes p<0.01; Chi square test (B); two-tailed Student’s t-test (F).

#### After-hyperpolarization (AHP) Analysis

Analysis of the AHP was performed on recordings where the cell was held at −80 mV and injected with a series of escalating 1-second current pulses. The AHP was analyzed using the first spike from the first trace where there was no greater than a 50 ms delay to the spike from current injection onset. Reported AHP values reflect the membrane potential at specified time intervals after the peak of the spike.

#### Pharmacology

In some electrophysiology recordings, mibefradil dihydrochloride hydrate (cat. no. M5441, Sigma Aldrich) was used at a concentration of 4 µM to fully inhibit Ca_V_3 family T-type Ca^2+^ channels (McDonough & Bean, 1998), and iberiotoxin (cat. no. STI-400, Alomone Labs) was used at 200 nM to fully inhibit BK channels.

### Tissue immunohistochemistry

Mice were anesthetized with isoflurane and brains were removed, fixed in 1% paraformaldehyde for 1 h, immersed in 30% sucrose in PBS and sectioned on a CM1850 cryostat (Leica). 14 µm parasagittal sections were processed for immunohistochemistry as described previously (Dell’Orco et al., 2015). Sections were kept stored at −80°C prior to antibody staining and imaging.

For molecular layer thickness measurements, Purkinje neurons were labelled with mouse anti-calbindin (1:1000, cat. no. C9848, Sigma-Aldrich) and goat anti-mouse Alexa488 conjugated secondary antibody (1:200, ref. no. A11001, Life Technologies Invitrogen). Sections were imaged using an Axioskop 2 plus microscope (Zeiss) at either 10x or 20x magnification. Measurements were performed using cellSens Standard image analysis software (Olympus). To measure molecular layer thickness in the anterior cerebellum, a line was drawn that measured 100 µm from the depth of the primary fissure along the length of the fissure, and the distance between the end of this line and the nearest Purkinje neuron cell body in lobule V was reported as the molecular layer thickness. To measure molecular layer thickness in the nodular zone, a line was drawn that measured 50 µm from the depth of the lobule IX/X fissure, and the distance between the end of this line and the nearest Purkinje neuron cell body in lobule IX or X was reported as the molecular layer thickness. Measurements of molecular layer thickness were performed in two sections per animal, and the reported molecular layer thickness for each animal is the mean of these two measurements. Sample preparation, imaging, and measurement for molecular layer thickness was performed with experimenter blind to genotype and treatment.

For double immunofluorescence experiments, Cic was labelled with rabbit anti-Capicua/CIC antibody (1:1000, cat. no. ab123822, Abcam) and goat anti-rabbit AlexaFluor 488 conjugated secondary antibody (1:200, cat. no. A11008, Life Technologies Invitrogen), while Purkinje cells were labelled with mouse anti-Calbindin antibody (1:1000, cat. no. C9848, Sigma-Aldrich) and goat anti-mouse AlexaFluor 594 (1:200, cat. no. A11005, Life Technologies Invitrogen). To quantify the intensity of Cic, images were analyzed using Plot Profile in ImageJ. To do this, a line was drawn across the Cic-positive area in the soma and the area under the curve was measured, and the somatic Cic intensity for that cell was defined as the area under the curve divided by the length of the line. Somatic Cic intensity was measured in ∼6 cells from both the anterior or nodular zone for each animal, and the mean of these values from the anterior cerebellum and nodular zone was determined for each animal. Those mean intensities were then all normalized to the average Cic intensity values from the anterior cerebellum of all wild-type animals to produce the values reported in Figure 4C. Sample preparation, imaging, and analysis was performed with experimenter blind to genotype.

### RNA sequencing and analysis of ion channel gene expression

#### RNA sequencing

Cerebella from 6-week-old ATXN2[Q127] and wild-type littermates (16 animals in each group) or 8-week-old ATXN2-BAC-Q72 and wild-type littermates (4 animals in each group) were used for RNA sequence analyses. Total RNA was isolated using miRNeasy Mini Kit (Qiagen Inc., USA) according to the manufacturer’s protocol. RNA quality was determined using the Bioanalyzer 2100 Pico Chip (Agilent). Samples with an RNA integrity number (RIN) >8 were used for library preparation using Illumina TrueSeq Stranded Total RNA Sample Prep with Ribo-Zero rRNA Removal Kit for mouse. Single-end 50-bp reads were generated on a Hiseq 2000 sequencing machine at the University of Utah Microarray and Genomic Analysis Shared Resource using Illumina Version 4 flow cells. Reads were then aligned to the mouse reference genome (mm10) by Novoalign (http://www.novocraft.com) and differentially expressed genes were identified as previously described (Dansithong et al., 2015).

#### Analysis of ion channel gene expression from SCA1 and SCA2 mice

Published data tables from a previous RNA sequencing analysis of gene expression analysis in ATXN1[82Q] mice (Ingram et al., 2016) and from a microarray analysis of gene expression in *Atxn1*^*154Q/2Q*^ mice (Gatchel et al., 2008) were downloaded from the NCBI Gene Expression Omnibus (accession number GSE75778 and GSE991, respectively). Data tables for ATXN2[Q127] mice and ATXN2-BAC-Q72 mice were generated as described above. From these data tables, genes that were found to show statistically significant differential expression in the original analysis for each dataset were selected for further analysis, along with corresponding fold change expression and log_2_ conversion of the fold change expression. In Figure 2B, the log_2_ conversion of the fold change expression relative to each model’s respective wild-type control is plotted.

Data tables of dysregulated genes were analyzed to identify shared transcripts across models using a custom script, which was executed using the Jupyter Notebook web application and running the Anaconda distribution of Python 3.6.3. This script and input data tables required to perform the analysis are available at https://github.com/chopravi/DiffEx_SCA1_SCA2. For each model, dysregulated channel genes were identified from the full list of differentially expressed genes in that model by searching the differentially expressed genes against the 270 genes found in mice from the IUPHAR classification for pore-forming (alpha subunit) ion channels (Alexander et al., 2019). The lists of dysregulated channel genes in each model were compared to identify all possible overlaps between models, and these overlaps are represented in Figure 2A using an UpSet plot (Lex, Gehlenborg, Strobelt, Vuillemot, & Pfister, 2014) along with an unscaled Venn diagram in Supplementary Figure 2A. For Figure 2A, broken axes were created in Illustrator (Adobe) using the direct output of the custom script referenced above.

To determine whether there was a statistically significant number of dysregulated channel genes shared across models, we tested the null hypothesis that the number of channel genes shared between these models could occur by chance, given the number of channel genes that are differentially expressed in each model. Specifically, to evaluate the statistical significance of the channels dysregulated in any 3 models, we tested the null hypothesis that 12 or more ion channel genes could be obtained from the union of the overlaps between any 3 models by chance. The probability for this overlap was determined by performing 1,000 simulations in which the appropriate number of channels were randomly sampled for each model from the IUPHAR database of all ion channels found in mice, at which point the number of shared channels in appropriate overlaps was counted. The null hypothesis was rejected and the number of channel genes that are dysregulated was deemed to be statistically significant if P<0.05.

### Gene expression in anterior cerebellum and nodular zone by qRT-PCR

#### Isolation of anterior cerebellum and nodular zone

Mice were euthanized following anesthesia with isoflurane, and cerebella were removed and placed in ice-cold PBS. The cerebellar hemispheres were removed with a razor blade, and the vermis was then laid on its lateral aspect and the lobules were visualized under a dissecting scope. The anterior cerebellum and nodular zone were separated from the remaining cerebellar tissue using glass microelectrodes pulled for patch clamp recordings. Once tissue was isolated, it was flash-frozen in liquid nitrogen and stored at −80°C until the time of processing.

#### RNA Extraction, cDNA synthesis, and quantitative real-time PCR

Total RNA from each harvested portion of mouse cerebellum was extracted using Trizol Reagent (Invitrogen) and subsequently purified using the RNeasy Mini Kit (Qiagen Inc., USA) following the manufacturer’s instructions. cDNA was synthesized from 1 μg of purified RNA using the iScript cDNA synthesis kit (Cat. no. 1708891, Bio-Rad). Quantitative real-time PCR assays were performed using the iQ SYBR Green Supermix (Cat. no. 1708880, Bio-Rad) in a MyiQ Single Color Real-Time PCR Detection System (Bio-Rad), with each reaction performed at a 20 μl sample volume in an iCycler iQ PCR 96-well Plate (Bio-Rad) sealed with Microseal optical sealing tape (Bio-Rad). The relative amount of transcript mRNA was determined using the comparative *C*_t_ method for quantitation (Livak & Schmittgen, 2001) with *Actb* mRNA serving as the reference gene. *C*_t_ values for each sample were obtained in triplicate and averaged for statistical comparisons. The primers used for qRT-PCR are listed below:

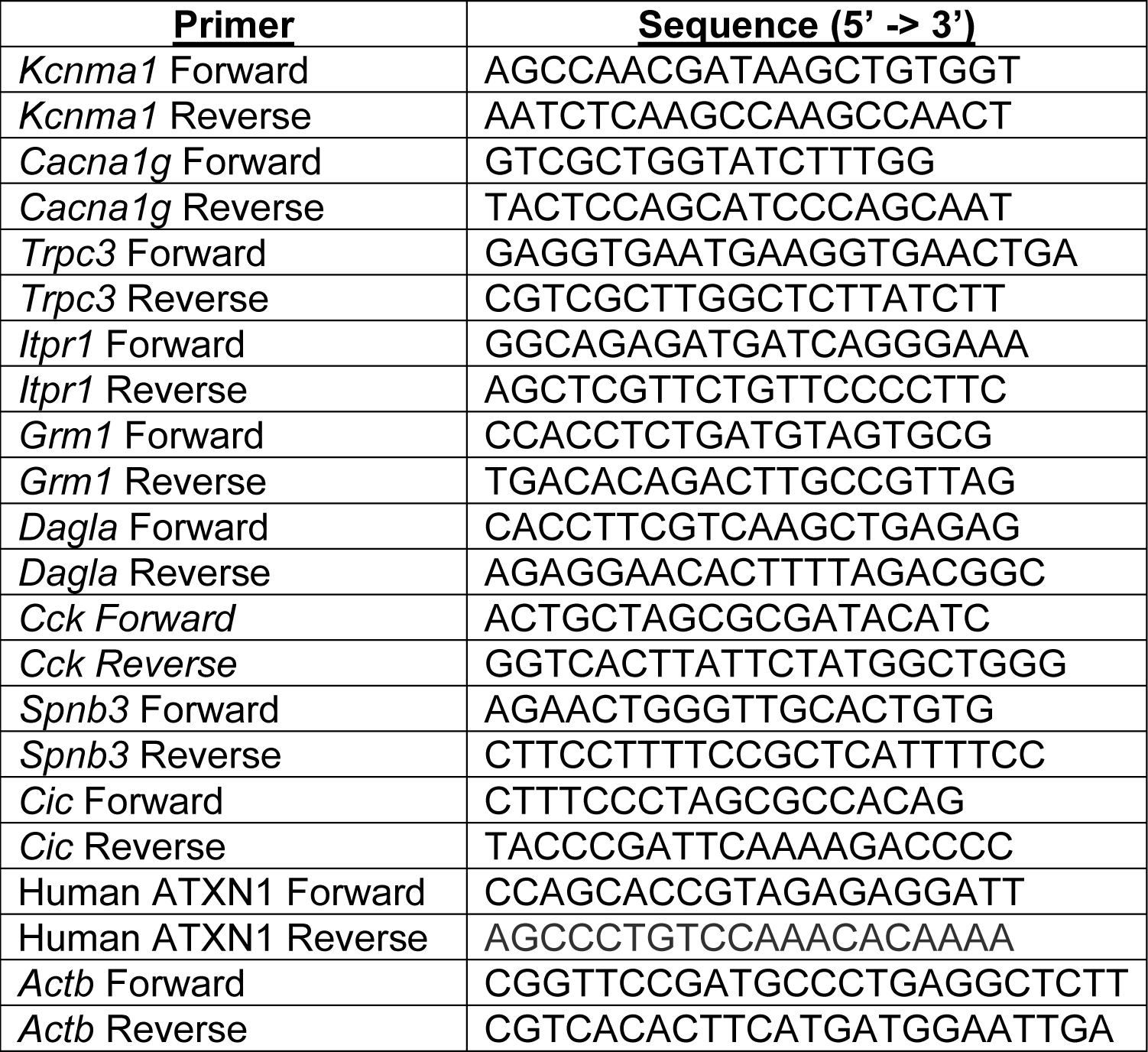

### Fast chromatin immunoprecipitation (qCHIP)

qCHIP was performed as previously described (Yellajoshyula et al., 2017) with minor modifications. A pool of isolated cerebellum tissue from P14 mice (three to six mice per genotype) was digested for 20 minutes with Pappain (SKU no. PAP, Brainbits) + DNase I (10 μg/ml; Sigma) followed by 10 fold dilution in PBS supplemented with 1 % FBS (fetal bovine serum). After centrifugation (300g, 5 minutes) the cell pellet was washed three times in PBS + 1% FBS and the resultant final pellet (containing a single cell suspension) was used to isolate chromatin. This suspension was sonicated until chromatin was sheared to a length of 200-500 bp. A quantity of sheared chromatin corresponding to one cerebellum from the chromatin pool was incubated with 50 μL of Rabbit anti-Atxn1 (kindly provided by Harry Orr), 2.5 μg Rabbit anti-Cap (cat. no. ab123822, Abcam), or 2.5 μg normalized Rabbit IgG (Santacruz) using Dynabeads (Invitrogen). After washing, elution and cross-link reversal, DNA from each ChIP sample and the corresponding input sample was purified.

Quantitative PCR (qPCR) was used to analyze the percentage of DNA recovered after ChIP from each sample. Each ChIP sample and a range of dilutions of the corresponding input sample (0.01 – 2% input) were quantitatively analyzed with gene-specific primers using the 7500 detection system (ABI) and SYBR qPCR Premix (Clontech). For each panel included in Figure 4 or supplementary Figure 4, the data presented are from an individual tissue preparation and immunoprecipitation. The primers used for qPCR are listed below:

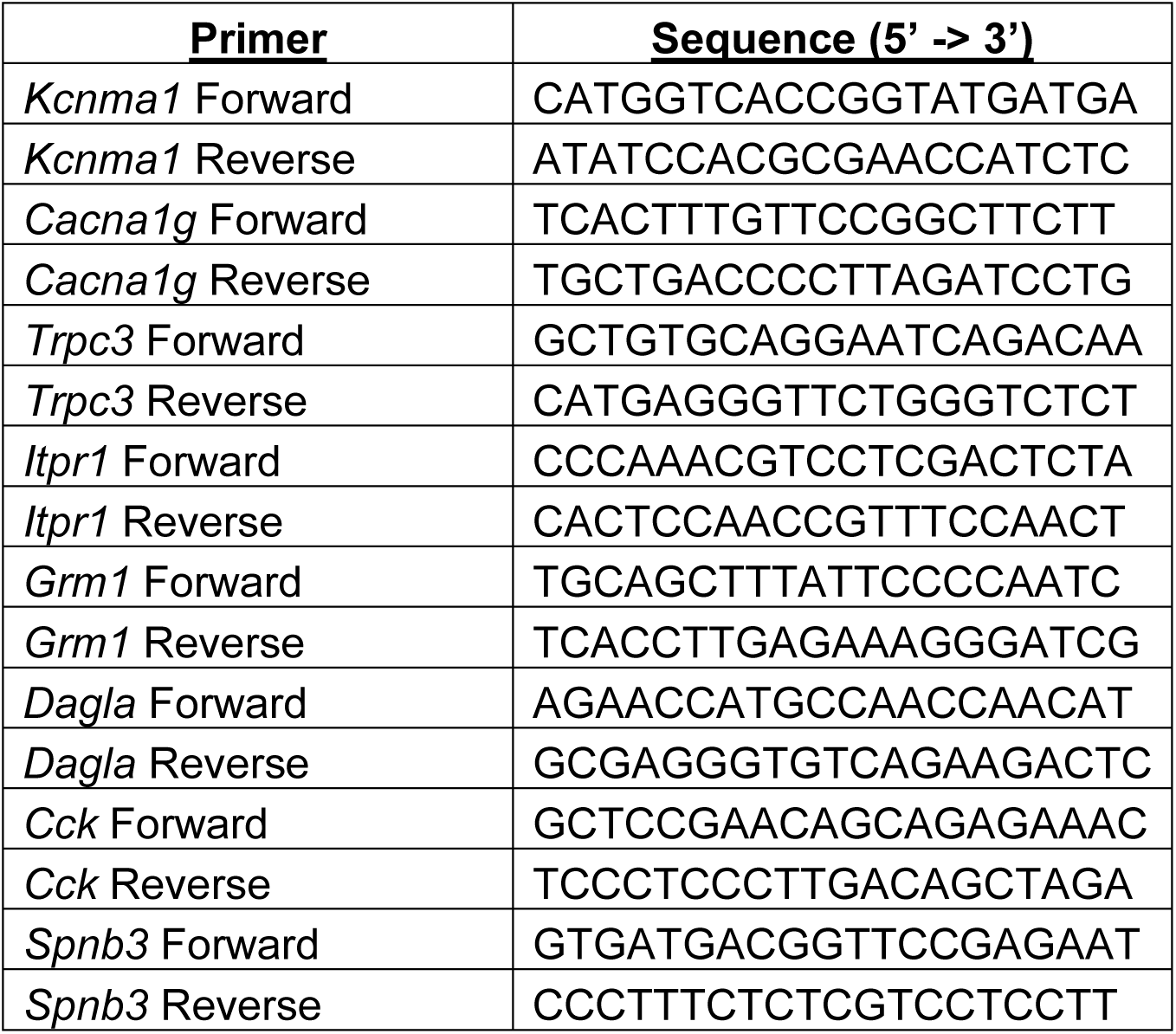

### Lentiviral transduction of Purkinje neurons

#### Viral vectors

Viral vectors were generated by and purchased from the University of Minnesota Viral Vector and Cloning Core (Minneapolis, MN). cDNA encoding either Capicua (Cic) or mCherry was packaged into a lentiviral vector under control of the human synapsin (hSyn) promoter. Myc-CICf was a gift from Huda Zoghbi (Addgene plasmid #48185; http://n2t.net/addgene:48185; RRID:Addgene_48185) (Kim, Lu, Zoghbi, & Song, 2013). Lentiviral titers were as follows: hSyn.Cic, 3.34×10^7^ viral particles/mL; hSyn.mCherry, 4.67×10^7^ viral particles/mL. hSyn.mCherry lentivirus was diluted to 3.33×10^7^ viral particles/mL in sterile phosphate buffered saline to facilitate equal delivery of volume and viral load into the cerebellar cortex.

#### Cerebellar delivery of lentivirus

Lentiviral vectors were delivered to the cerebellar cortex using stereotaxic surgical techniques. ATXN1[82Q] or wild-type controls at P28-P38 were used for surgery. Mice were anesthetized under 5% isoflurane inhalation and were maintained at 1.5% isoflurane for the duration of the procedure. A single craniotomy above the nodular zone was performed with a small drill burr at bregma −7.0 mm in the anterior/posterior plane. 3.0 µL lentivirus was loaded into a 10 µL Hamilton syringe (BD Biosciences, San Jose, CA), which was then slowly lowered into the nodular zone to a depth of bregma −3.8 mm in the dorsal/ventral plane. A series of pilot injections using 3.0 µL of hSyn.mCherry lentivirus was used to confirm the surgical coordinates. For electrophysiological recordings and molecular layer thickness measurements, mice were injected with either 3.5 µL hSyn.mCherry (control) or a combination of 3.0 µL hSyn.Cic + 0.5 µL hSyn.mCherry for verification of infection in lobule IX. Lentivirus was delivered at a rate of 500 nL/minute using an injection pump (UMC4, World Precision Instruments, Inc., Sarasota, FL). After completion of delivery, the syringe was allowed to sit for two minutes before slight retraction, after which the syringe was left in place for six additional minutes to prevent backflow of the lentivirus into the syringe tract. After removal of the syringe, the scalp was sutured, and the mouse was monitored during recovery from anesthesia. Post-operative care was provided for at least seven days after surgery, and carprofen (2.5 mg/kg subcutaneous) was administered for the first 72 hours after surgery.

### Chemicals

Reagents and chemicals were obtained from Sigma-Aldrich unless otherwise specified.

### Statistical Analysis

Statistical tests are described in the Figure legends for all data. Data are expressed as mean ± SEM unless otherwise specified. Sample size is included in each figure panel, either by plotting of individual data points or by including the number of cells (n) within the figure panel. Studies were powered and analysis was performed assuming unequal variance between groups. Data were analyzed using GraphPad Prism (GraphPad) and Excel (Microsoft)

## Results

### Purkinje neurons from the anterior cerebellum are more vulnerable than Purkinje neurons from the nodular zone in SCA1 mice

We began by quantifying the kinetics of neurodegeneration in Purkinje neurons from anterior cerebellum and compared them to Purkinje neurons from the nodular zone (Figure 1A), as previous studies have noted relative sparing of Purkinje neurons in lobule X found within the nodular zone (Clark et al., 1997). Measurement of membrane capacitance (Figure 1B-1C) and molecular layer thickness (Figure 1D-1E) showed that dendritic degeneration begins at P35 in the Purkinje neurons from the anterior cerebellum of ATXN1[82Q] mice compared to P105 in the nodular zone. The early onset of Purkinje neuron degeneration starting at P35 is not explained by differences in ATXN1[82Q] transgene expression at that age (Supplementary Figure 1E), though we found reduced transgene expression at P105 (Supplementary Figure 1F). These findings suggest that Purkinje neurons from the anterior cerebellum are more vulnerable in ATXN1[82Q] mice and argue for the existence of a disease pathway unique to anterior Purkinje neurons in the ATXN1[82Q] model of SCA1.

### A functionally related module of channels is dysregulated uniquely in the anterior cerebellum of SCA1 mice

Polyglutamine-expanded ATXN1 is thought to exert its pathogenic effect in Purkinje neurons primarily by disrupting gene expression. Dysregulation of individual transcripts and transcript modules has been suggested as the root cause for disease (Paulson, Shakkottai, Clark, & Orr, 2017), and unbiased analysis of ataxia genes demonstrates that these genes fit within a limited number of pathways to unify Purkinje neuron vulnerability across spinocerebellar ataxias (Bettencourt et al., 2014). Altered Purkinje neuron intrinsic excitability has emerged as one particularly important pathway (Bushart & Shakkottai, 2019), suggesting that there may be a limited number of channels that unify Purkinje neuron dysfunction across ataxias.

To identify those ion channel genes whose dysregulation was likely to reflect a biologically-relevant disease pathway for Purkinje neurons, we compared previously-published cerebellar whole-transcriptome gene expression datasets from models of spinocerebellar ataxias type 1 and 2, as Purkinje neurons are prominently affected in both conditions (Ferrer et al., 1994; Koeppen, 2005). Comparison of shared dysregulated transcripts using RNA sequencing or microarray studies from ATXN1[82Q] (Ingram et al., 2016), *Atxn1*^*154Q/2Q*^ (Gatchel et al., 2008), ATXN2[Q127], and ATXN2-BAC-Q72 models (Dansithong et al., 2015) revealed twelve pore-forming (alpha subunit) ion channel genes that are differentially expressed in at least three of the four models (Figure 2A-B, Supplementary Figure 2A). Among these twelve genes, there are four genes for which mutations are known to cause human ataxia syndromes, namely *Kcnma1, Cacna1g, Trpc3*, and *Kcnc3* (Coutelier et al., 2015; Fogel, Hanson, & Becker, 2016; Liang et al., 2019; Waters et al., 2006). This finding suggests that dysregulation of these channel genes specifically is likely to be a root cause for Purkinje neuron pathology in cerebellar ataxia.

Downregulation of *Kcnma1*-encoded large-conductance calcium-activated K^+^ (BK) channels has been validated experimentally to be important for the pathogenesis of both SCA1 and SCA2 (Chopra, Bushart, et al., 2018; Dell’Orco, Pulst, & Shakkottai, 2017; Dell’Orco et al., 2015). However, in wild-type neurons, the severe Purkinje neuron dysfunction observed in models of SCA1 or SCA2 cannot be recapitulated by pharmacologic blockade of BK (Sausbier et al., 2004), suggesting that Purkinje neuron dysfunction in SCA1 and SCA2 must also involve other channels. Importantly, such severe dysfunction is readily apparent with intracellular Ca^2+^ chelation (Edgerton & Reinhart, 2003), and recent work in a model of spinocerebellar ataxia type 7 has demonstrated that dysregulation of both BK channel expression and Ca^2+^ homeostasis promote Purkinje neuron degeneration (Stoyas et al., 2020). Given that our analysis of SCA1 and SCA2 models reveals dysregulation of not just *Kcnma1*, but also the ataxia-linked Ca^2+^ channel genes *Cacna1g* and *Trpc3*, we hypothesized that severe Purkinje neuron dysfunction observed in SCA1 might arise due to synergistic dysfunction of an ion channel module that contains BK and Ca^2+^ sources (Figure 2C).

To investigate whether coordinate dysregulation of this module may explain accelerated degeneration of Purkinje neurons from the anterior cerebellum, we performed qRT-PCR using macrodissected anterior cerebellum and nodular zone and investigated expression of a panel of genes implicated in SCA1 pathogenesis. These genes included the ion channel module described above (*Kcnma1, Cacna1g*, and *Trpc3*) as well as *Cck* (Ingram et al., 2016), *Dagla, Prkcg* (Chopra, Wasserman, Pulst, De Zeeuw, & Shakkottai, 2018), *Itpr1* (Lin, Antalffy, Kang, Orr, & Zoghbi, 2000), and *Grm1* (Shuvaev, Hosoi, Sato, Yanagihara, & Hirai, 2017). At P35, when neurodegeneration is evident only in the anterior cerebellum (see Figure 1), *Kcnma1, Cacna1g, Trpc3, Itpr1*, and *Grm1* were downregulated only in the anterior cerebellum. In contrast, *Cck, Dagla*, and *Prkcg* were downregulated in both regions (Figure 2E-2F). Once neurodegeneration has also begun in the nodular zone (P105, see Figure 1), *Kcnma1, Cacna1g, Itrp1*, and *Grm1* were also downregulated in the nodular zone (Figure 2G-2H). Taken together, these findings demonstrate tight spatiotemporal correlation between downregulation of this ion channel module and the onset of Purkinje neuron degeneration. We identify the critical genes in this module to be *Itpr1, Cacna1g*, and *Kcnma1* and support a potential role for *Grm1*.

### Regionally-dysregulated ion channel genes form a functional module critical for Purkinje neuron pacemaking

To determine whether regional downregulation of ion channel genes correlates with abnormal Purkinje neuron physiology, acute slice patch clamp recordings were performed in Purkinje neurons from the anterior cerebellum and nodular zone in ATXN1[82Q] and wild-type mice at P35. As previously reported, Purkinje neurons from ATXN1[82Q] mice are hyperexcitable and lack normal pacemaking at P35, with disrupted pacemaking arising from depolarization block (Bushart et al., 2018; Dell’Orco et al., 2015). We confirmed these prior findings of depolarization block in up to 50% of Purkinje neurons in the anterior cerebellum. Surprisingly, there were no differences in firing between wild-type and ATXN1[82Q] Purkinje neurons in the nodular zone (Figure 3A-3C). Additional analysis of firing properties from all firing cells, namely frequency (Supplementary Figure 3A) and coefficient of variation (Supplementary Figure 3B) also revealed no difference in the nodular zone. A reduction in the afterhyperpolarization (AHP) has been suggested as the cause for depolarization block in ATXN1[82Q] Purkinje neurons (Bushart et al., 2018; Dell’Orco et al., 2015), and there was a robust reduction in the AHP in ATXN1[82Q] Purkinje neurons from the anterior cerebellum (Figure 3D) but not the nodular zone (Figure 3E). Taken together, these findings demonstrate that there is Purkinje neuron hyperexcitability and dysfunction only in the anterior cerebellum at P35, consistent with the regionally-restricted downregulation of *Kcnma1* and its potential Ca^2+^ sources.

Canonically, P/Q-type Ca^2+^ channels are thought of as the primary Ca^2+^ source that recruits calcium-activated K^+^ channels during the Purkinje neuron action potential (Womack, Chevez, & Khodakhah, 2004). However, the proposed ion channel does not include P/Q-type channels but does include other Ca^2+^ channels, namely the T-type Ca^2+^ channel Ca_V_3.1 (encoded by *Cacna1g*) and the type 1 inositol 1,4,5-triphosphate (IP3) receptor (encoded by *Itpr1*). Pharmacologic ion channel blockade was utilized in wild-type Purkinje neurons to determine whether these channels operate as upstream Ca^2+^ sources for BK in its role as a regulator of Purkinje neuron pacemaking. Consistent with prior studies (Edgerton & Reinhart, 2003; Sausbier et al., 2004), isolated blockade of BK with saturating concentrations of iberiotoxin was not sufficient to produce depolarization block (Figure 3F-3G). However, when BK channels were blocked along with T-type Ca^2+^ channels (using mibefradil at a dose specific for T-type channels in Purkinje neurons (McDonough & Bean, 1998)), wild-type Purkinje neurons became irregular or undergo depolarization block (Figure 3F, 3H). These data suggest that downregulation of BK along with one or more of these Ca^2+^ sources could be sufficient to explain the observed depolarization block, and also demonstrate that this group of ion channel genes comprises a functional module critical for Purkinje neuron pacemaking.

### Higher Cic expression in Purkinje neurons from the anterior cerebellum of SCA1 mice is associated with repression of key ion channel genes

Having demonstrated the critical role for this ion channel module in Purkinje neurons, and knowing that its downregulation is tightly correlated with Purkinje neuron degeneration, we set out to explore the mechanism by which polyglutamine-expanded ATXN1 downregulates this ion channel module. We focused specifically on the transcriptional repressor Cic, which forms a native complex with ATXN1 that is central to SCA1 pathogenesis (Fryer et al., 2011; Lam et al., 2006). Regional analysis of Cic mRNA (Figure 4A) revealed that *Cic* expression is 28.6% higher in the anterior cerebellum at P35 (ATXN1[82Q]: 1.286±0.067, wild-type: 1.026±0.071). Previous reports have suggested that cerebellar Cic expression is largely restricted to immature granule cells (Lee et al., 2002), so the cell-type specificity of this effect was confirmed by immunohistochemistry, which showed an 18.0% higher nuclear Cic signal in anterior cerebellar Purkinje neurons from ATXN1[82Q] mice (Figure 4B-4C) P35 (ATXN1[82Q]: 1.180±0.026, wild-type: 1.000±0.047). These findings demonstrate that modestly greater Cic expression in the anterior cerebellum correlates with dysregulated ion channel gene expression and Purkinje neuron degeneration in this region of the cerebellum in ATXN1[82Q] mice.

Because Cic functions principally as a transcriptional repressor, we explored whether Cic binding to ion channel genes is uniquely increased in the presence of polyglutamine-expanded ATXN1. Using the *Atxn1*^*154Q/2Q*^ model of SCA1, where expression of Atxn1 is at endogenous levels (Watase et al., 2002), we found that while Cic binding for all genes was above IgG background, there was no polyglutamine length-dependent change in promoter occupancy (Figure 4D-4E, Supplementary Figure 4A-4C). In contrast, Atxn1 showed robust polyglutamine length-dependent increases in baseline binding to all SCA1-associated genes (Figure 4F-4G, Supplementary Figure 4D-4G). Taken together, these findings do not support the hypothesis that decreased ion channel expression is explained by increased binding of Cic specifically to ion channel genes, and suggest instead that ion channel gene expression may be sensitive to changes in Cic protein levels.

### Increased Cic expression results in Purkinje neuron hyperexcitability and accelerates Purkinje neuron degeneration

To investigate the causal connection between regionally greater Cic expression and ion channel dysregulation, we utilized a lentiviral vector to increase Cic expression in the nodular zone. Lentiviral injection produces reliable overexpression of coinjected *mCherry* within the posterior half of lobule IX (Figure 5A), so patch clamp recordings were performed in Purkinje neurons from this area. At P35, a significant proportion of Purkinje neurons from nodular zone lobule IX were found to display irregular spiking or depolarization block (Figure 5B-5C) in ATXN1[82Q] mice injected with *Cic* lentivirus. Of note, lentiviral transduction by itself led to a greater degree of irregularity, as Purkinje neurons from an area that robustly expressed *mCherry* were more irregular than those in an area that was poorly transduced (Supplementary Figure 5A). Despite this, there was still a clear effect of Cic expression on the degree of irregularity (Supplementary Figure 5B). Cic-dependent disruptions in Purkinje neuron pacemaking occurred in association with a reduction in the AHP (Figure 5D). These findings demonstrate that increasing Cic levels makes ATXN1[82Q] Purkinje neurons hyperexcitable.

Next, we investigated whether increased expression of Cic in the nodular zone impacted the kinetics of Purkinje neuron degeneration. Measurement of molecular layer thickness in the posterior half of lobule IX revealed accelerated Purkinje neuron degeneration in ATXN1[82Q] mice injected with *Cic* lentivirus (Figure 5E-5F). These data demonstrate that greater expression of Cic and Cic-dependent hyperexcitability is likely to be responsible for accelerated degeneration of anterior cerebellar Purkinje neurons in ATXN1[82Q] mice.

## Discussion

Selective neuronal vulnerability is a universal feature of neurodegenerative disease, but the relevant pathways that differentiate affected cell types from protected cell types are not well understood. By examining subpopulations within a prominently affected cell type (Purkinje neurons), we identified a key disease pathway for the pathogenesis of SCA1. These findings not only provide important insights into the pathobiology of spinocerebellar ataxia, but also validate the importance of cell subtype-level analysis as a platform for pathway discovery in neurodegenerative disease.

### Regional patterns of vulnerability in cerebellar degeneration

The current study not only confirms prior observations of an antero-posterior gradient of Purkinje neuron pathology in this model of SCA1 (Clark et al., 1997), but adds to a robust literature exploring this phenomenon more widely in cerebellar ataxia syndromes. Human imaging and autopsy studies have demonstrated that the posterior cerebellum is generally more resistant to degeneration (Jung et al., 2012; Kume, Takahashi, Hashizume, & Asai, 1991), and animal studies have suggested that this may stem from intrinsic differences in the resilience of Purkinje neurons across the antero-posterior axis, including differences in basal expression of genes that modulate protein homeostasis (Chung et al., 2016) or inflammation (Martin et al., 2019). Our findings support a different hypothesis for SCA1 that may be a general feature of degenerative cerebellar ataxias, namely, rather than simply being more vulnerable to stress, anterior Purkinje neurons have a different regulatory program for key cellular functions (intrinsic excitability and its control by Cic in SCA1) that can be acted upon by relevant pathogenic proteins (i.e. polyglutamine-expanded ATXN1).

Several whole-transcriptome expression studies support the idea that tissue-specific gene expression networks can form a substrate for neurodegeneration, including in spinocerebellar ataxias broadly (Bettencourt et al., 2014) and in this model of SCA1 specifically (Ingram et al., 2016). In our study, a more limited panel of genes is explored, but there is nevertheless a clear “module” of ion channel genes that together mediate a specific cellular function and are coordinately regulated. Furthermore, our study suggests that Cic is the regulator for this module, which not only establishes a novel role for Cic as a regulator of excitability in Purkinje neurons but also provides a mechanistic link between polyglutamine-expanded ATXN1 and abnormal Purkinje neuron physiology in SCA1.

The current study did not identify the mechanism underlying greater Cic expression in the anterior cerebellum of ATXN1[82Q] mice, though it demonstrated that the effect is mediated through higher *Cic* mRNA levels. Canonically, Cic function is regulated through control of Cic protein abundance, with steady-state levels set through a balance between binding to stabilizing partners (including ATXN-1 (Lam et al., 2006)) and receptor tyrosine kinase-dependent phosphorylation and degradation (Jiménez, Shvartsman, & Paroush, 2012). Additional studies are required to determine the basis for our finding of increased Cic expression in anterior cerebellar Purkinje neurons, as this may represent a phenomenon relevant to human disease.

Notably, our findings indicate that a relatively modest elevation of Cic (∼20% in the anterior cerebellum) is nevertheless sufficient to produce Purkinje neuron hyperexcitability and degeneration. This complements recent findings in human patients with heterozygous truncation mutation in CIC, which led to a neurobehavioral disorder that was likely developmental in origin (Lu et al., 2017). It is clear that neurons across a variety of brain regions are very sensitive to CIC gene dosing, and that CIC levels are likely to be tightly regulated throughout the lifespan. Because this regulation appears to break down in SCA1, treatments that target regulators of CIC (or target CIC itself) to modestly lower expression could produce profound symptomatic and neuroprotective benefits in patients with SCA1.

### Purkinje neuron excitability and Cic in SCA1 pathogenesis

Cic forms a native complex with ATXN1 that is essential for SCA1 pathogenesis (Fryer et al., 2011; Lam et al., 2006; Rousseaux et al., 2018). While significant progress has been made in understanding the ATXN1/Cic complex, including a validated partial crystal structure (Kim et al., 2013), much less is known about the mechanisms by which this complex drives Purkinje neuron degeneration. Our study suggests that Purkinje neuron excitability may be one such mechanism, and specifically highlights a functionally related group of channels (*Cacna1g, Itpr1*, and *Kcnma1*) that we also show to be critical for Purkinje neuron pacemaking.

Abnormal Purkinje neuron excitability has emerged as an important pathogenic mechanism across multiple polyglutamine ataxias (Bushart & Shakkottai, 2019; Paulson et al., 2017). Disrupted Purkinje neuron pacemaking in particular has been found in models of SCA1 (Dell’Orco et al., 2015; Hourez et al., 2011), SCA2 (Dell’Orco et al., 2017; Kasumu et al., 2012), SCA3 (Shakkottai et al., 2011), SCA6 (Jayabal, Chang, Cullen, & Watt, 2016), and SCA7 (Stoyas et al., 2020), with many studies demonstrating that treatments which improve pacemaking also slow Purkinje neuron degeneration. Purkinje neuron pacemaking depends on a suite of ion channels whose biophysical properties and expression levels are precisely tuned to produce a regenerative firing cycle (Raman & Bean, 1999). Our study supports two important conclusions about this suite of channels and their dysregulation in SCA1 (Figure 6): 1) disrupted pacemaking in SCA1 occurs through downregulation of tightly-coupled channels with a synergistic impact on spiking, 2) the affected channels are under coordinate control through Cic.

**Figure 6.**
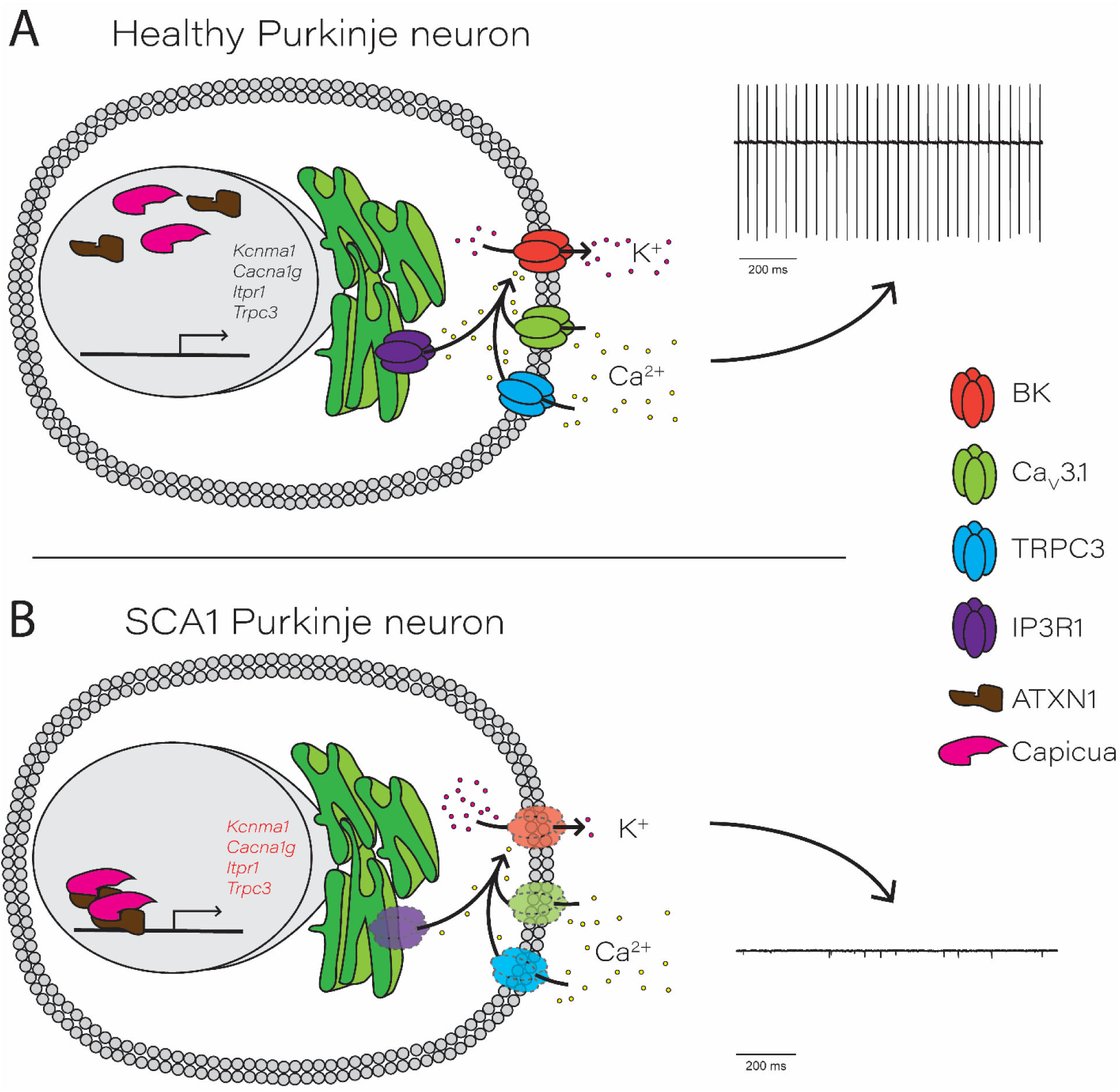
Proposed role for Capicua in transcriptional repression of essential Purkinje neuron ion channel genes. In healthy, wild-type Purkinje neurons, wild-type ATXN1 is located in both the cytosol and nucleus. Capicua, located in the nucleus, exerts basal transcriptional repression but allows transcription of key Purkinje neuron ion channel genes. Ca_v_3.1 and TRPC3, located on the plasma membrane, and IP3R1, located on the endoplasmic reticulum membrane, act as Ca^2+^ sources for the BK channel. BK channel activation allows for normal potassium efflux during the action potential, which supports pacemaker firing (right). (B) In SCA1, increased repression of key Purkinje neuron ion channel genes is driven by the interaction between Capicua and polyglutamine-expanded ATXN1. Transcriptional repression results in reduced expression of ion channel proteins, leading to decreased BK channel activity and an inability to support pacemaker firing.

In conclusion, our study identifies a likely Cic-dependent ion channel module whose critical role in Purkinje neuron pacemaking has not previously been described. The relevance of this module is likely to extend beyond SCA1, as our own analysis of whole-transcriptome data from SCA2 suggests that members of the module are dysregulated in SCA2, and a recent study in a model of SCA7 also demonstrates reduced transcript levels for all genes from this module (Stoyas et al., 2020). Given that ATXN2 and ATXN7 have not been described to act through Cic, it is tempting to speculate that the ion channel module we have identified represents an important shared pathway for Purkinje neuron pathology in many spinocerebellar ataxias. For this reason, our study in SCA1 highlights the potential for shared pathogenic processes with broad applicability to cerebellar disease.

## Acknowledgements

This work was supported by the NIH R01 NS085054 (V.G.S), T32GM007863 (R.C.) and the National Ataxia Foundation Research Post-Doctoral Fellowship Award (R.C.). We would like to thank Brandon Lee and Annie Zalon for technical support.

## Conflict of Interest Statement

Nothing to report.

**Supplementary Figure 1.**
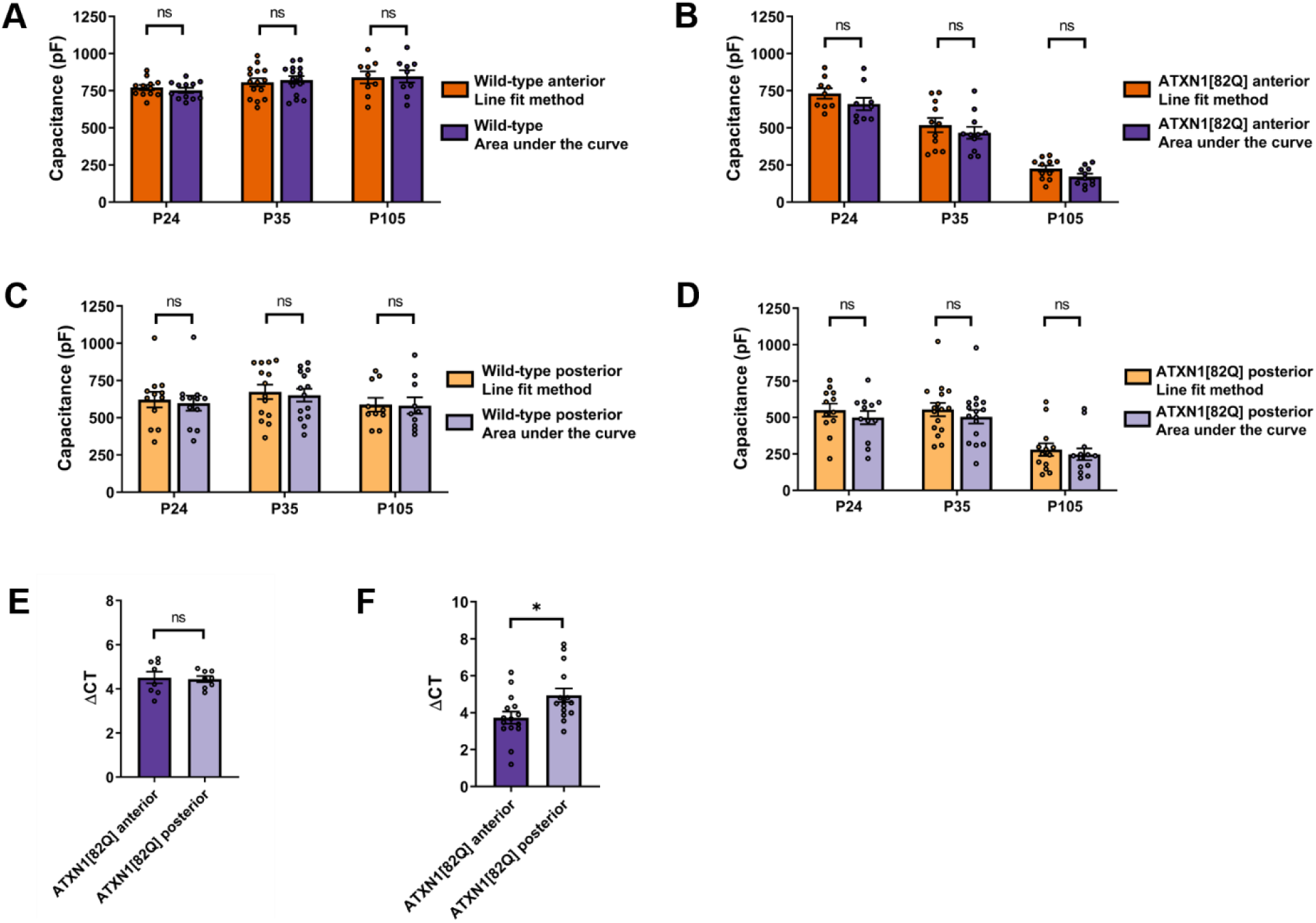
Detection of Purkinje neuron capacitance and cerebellar ATXN1[82Q] transgene expression. Capacitance data in Figure 1B-1C was fit using a two-exponential decay function based on a model for the Purkinje neuron as a two-compartment equivalent circuit (Llano et al., 1991). (A) Capacitance measurements from wild-type Purkinje neurons in the anterior cerebellum of Figure 1B were compared using the two-exponential decay function and an alternative method of measuring capacitance using area under the curve of a −10 mV voltage step from −70 mV to −80 mV. (B) Similar to (A), capacitance measurements are compared for ATXN1[82Q] Purkinje neurons in the anterior cerebellum of Figure 1B using the two-exponential decay function and the area under the curve method. (C) Similar to (A), capacitance measurements are compared for wild-type Purkinje neurons in the nodular zone of Figure 1C using the two-exponential decay function and area under the curve method. (D) Similar to (C), capacitance measurements are compared for ATXN1[82Q] Purkinje neurons in the nodular zone of Figure 1C using the two-exponential decay function and area under the curve method. (E) Relative human ATXN1[82Q] transgene expression is displayed for macrodissected anterior cerebellum and nodular zone from ATXN1[82Q] mice at P35. (F) Relative human ATXN1[82Q] transgene expression is displayed for macrodissected anterior cerebellum and nodular zone from ATXN1[82Q] mice at P105. * denotes p<0.05; ns denotes p>0.05; two-way repeated measures ANOVA with Holm-Sidak correction for multiple comparisons (A-D); two-tailed Student’s t-test (E-F).

**Supplementary Figure 2.**
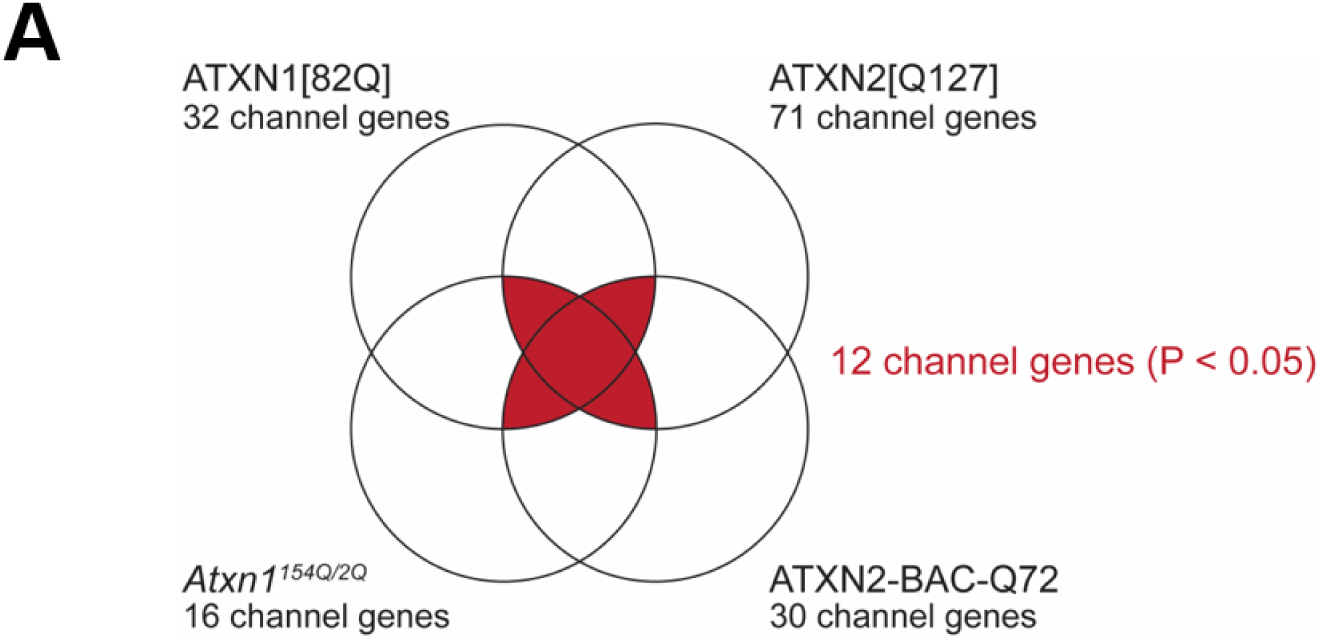
Ion channel transcripts showing dysregulated expression in mouse models of SCA1 and SCA2. (A) Dysregulated ion channel genes that are found to be shared across whole cerebellar gene expression datasets in SCA1 and SCA2 mouse models. The red overlap represents ion channel genes that are dysregulated in either three or four models. P-values reflect the likelihood of an equivalent number of channels (or more) being dysregulated in any three and in all four models by chance (see methods section).

**Supplementary Figure 3.**
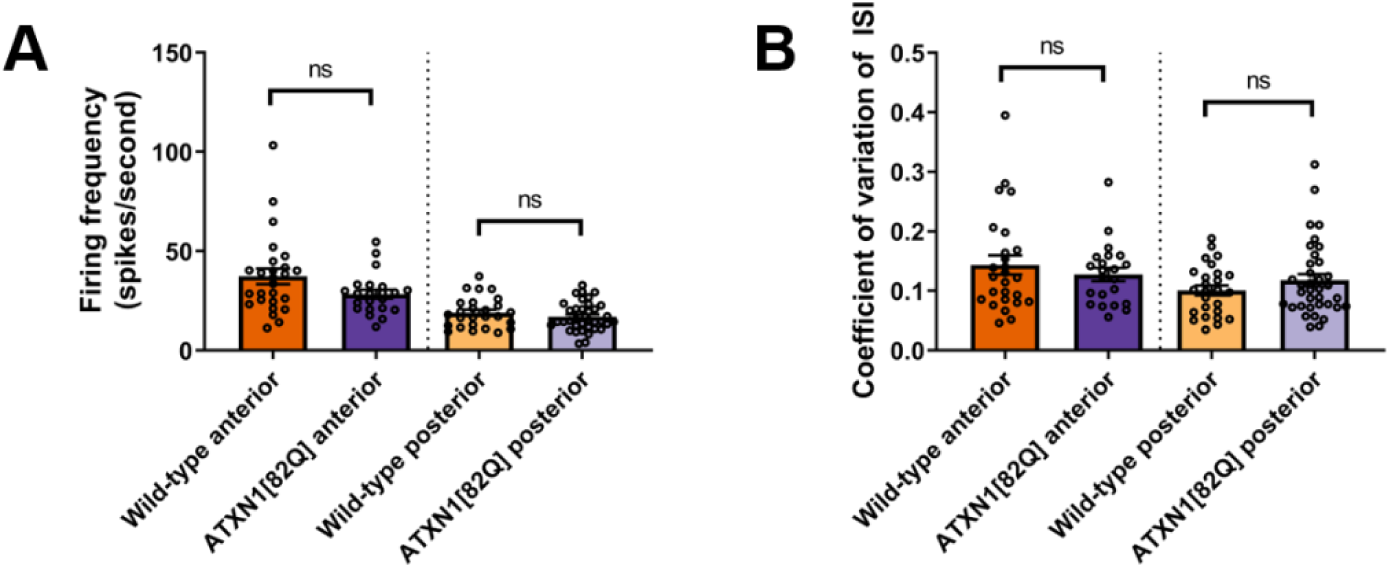
Spontaneous Purkinje neuron firing in ATXN1[82Q] mice and wild-type controls. Patch clamp electrophysiology in acute cerebellar slices from ATXN1[82Q] mice and wild-type controls was performed at P35. (A) Firing frequency in anterior cerebellum and the nodular zone is shown. (B) Coefficient of variation (CV) of the interspike interval (ISI) is shown for Purkinje neurons in the anterior cerebellum and nodular zone. ns denotes p>0.05; two-tailed Student’s t-test.

**Supplementary Figure 4.**
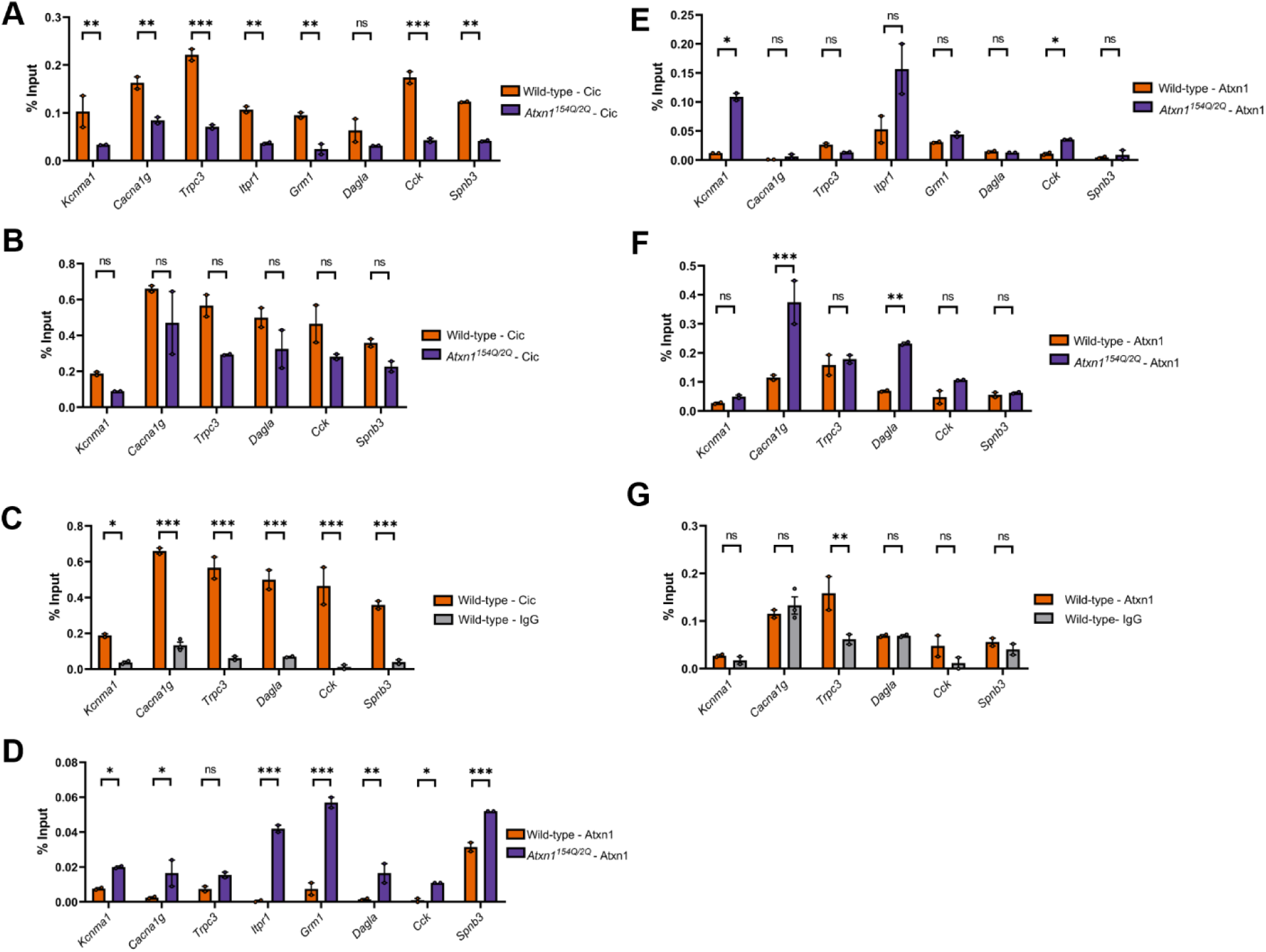
Enhanced binding of Atxn1 and Capicua to the promoter regions of ion channel module genes. (A-G) Quantitative Chromatin immunoprecipitation (qChIP) demonstrating the association of Capicua (Cic) and ATXN1 at the promoter of ion channel genes from sonicated chromatin derived from P14 whole cerebellar extracts. Binding, represented as % input (Y-axis) demonstrated for Cic (A-C) and ATXN1 (D-G) comparing their relative binding on ion channel genes in *Atxn1*^*154Q/2Q*^ mice and wild-type controls (A-B, D-F) and binding over background relative to their respective isotype control IgG (rabbit IgG) (C, G). * denotes p<0.05; ** denotes p<0.01; *** denotes p<0.001; ns denotes p>0.05; two-tailed Student’s t-test with Holm-Sidak correction for multiple comparisons.

**Supplementary Figure 5.**
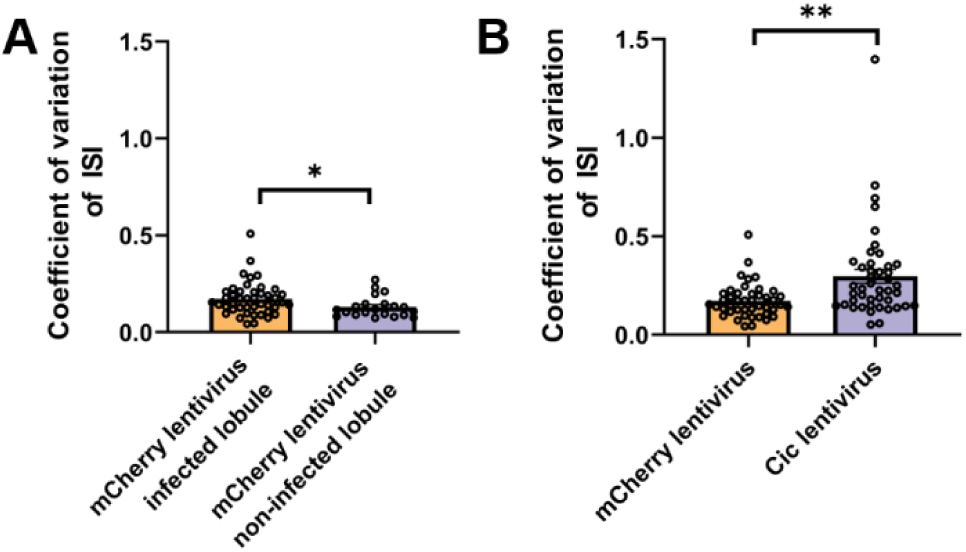
Spontaneous Purkinje neuron spiking after ectopic lentiviral expression of Capicua. Patch clamp electrophysiology in acute cerebellar slices was performed in the nodular zone of ATXN1[82Q] mice 10 days after lentivirus injection. (A) Coefficient of variation (CV) of the interspike interval (ISI) of Purkinje neuron spiking in the transduced area (lobule IX) and non-transduced area (lobule X) of mCherry lentivirus-injected ATXN1[82Q] mice. (B) CV of the ISI of Purkinje neuron spiking in the transduced area (lobule IX) of ATXN1[82Q] cerebellum for mice injected with either mCherry lentivirus or Cic lentivirus. * denotes p<0.05; ** denotes p<0.01; two-tailed Student’s t-test.

